# EEG Artifact to Signal: Predicting Horizontal Gaze Position from SOBI-DANS Identified Ocular Artifact Components

**DOI:** 10.1101/2020.08.29.272187

**Authors:** Rui Sun, Cynthia Chan, Janet Hsiao, Akaysha C. Tang

## Abstract

Ocular artifact in EEG has long been viewed as a problem for interpreting EEG data in basic and applied research. The removal of such artifacts has been an on-going effort over many decades. We have recently introduced a hybrid method combining second-order blind identification (SOBI) with DANS, a novel automatic identification method, to extract components containing specifically signals associated with horizontal and vertical saccadic eye movements (H and V Comps) and found that these components’ event-related potentials in response to saccadic eye movement are systematically modulated by movement directions and distances. Here in a case study, taking advantage of signals about gaze positions contained in the ocular artifact components, we introduced a novel concept of EEG-based virtual eye tracking (EVET) and presented its first prototype. Specifically, we determined (1) the amount of data needed for constructing models of horizontal gaze positions; (2) the asymptotic performance levels achieved with such models. We found that for the specific calibration task, 4 blocks of data (4 saccades per target position) are needed for reaching an asymptotic performance with a prediction accuracy of 0.44 and prediction reliability of 1.67. These results demonstrated that it is possible to track horizontal gaze position via EEG alone, ultimately enabling coregistration of eye movement and the neural signals.

## 1. Introduction

Electroencephalography (EEG) and EEG-based source imaging have been widely used in variety of basic, clinical, educational, and commercial neuroscience research and applications (Donchin, Spencer, & Wijesinghe, 2000; Niedermeyer & Lopes Da Silva, 2004; Khushaba, et al., 2013; Luck, 2014; McLoughlin, Makeig, & Tsuang, 2014; Zink, Hunyadi, Van Huffel, & De Vos, 2016; Dikker, et al., 2017; Cavanagh, 2018; He, Sohrabpour, Brown, & Liu, 2018; Protzak & Gramann, 2018; Lau-Zhu, Lau, & McLoughlin, 2019). While advances have been made in the development of new electrodes for better signal quality (Casson, 2019) and ease of application (Lopez-Gordo, Sanchez-Morillo, & Pelayo Valle, 2014; Kam, et al., 2019) and in the application of blind source separation algorithms (Bell & Sejnowski, 1995; Belouchrani, Abed-Meraim, Cardoso, & Moulines, 1997) for artifact removal (Jung, et al., 2000; Joyce, Gorodnitsky, & Kutas, 2004), (for review, see Bridwell, et al., 2018; Mannan, Kamran, & Jeong, 2018), the presence of the large amplitude artifact associated with eye movement continues to constrain the way EEG could be used to investigate the neural processes underlying natural cognition. For instance, in part to minimize the contamination of the EEG by ocular artifacts associated with eye movement, typical experimental paradigms require fixation of the eye at a particular location prior and during stimulus presentation. Consequently brain-behavior relations cannot be investigated in the natural context of free eye movement.

While a persistent effort has been made over the years to extract and identify the components associated with eye movements and other artifacts from the original EEG data, the general approach has been to use BSS to isolate the ocular artifact components, remove the artifactual components from the original EEG data, and then ultimately perform the analysis on the cleaned EEG data (for reviews, see Croft & Barry, 2000; Croft, Chandler, Barry, Cooper, & Clarke, 2005; Dimigen, Sommer, Hohlfeld, Jacobs, & Kliegl, 2011; Urigüen & Garcia-Zapirain, 2015; Islam, Rastegarnia, & Yang, 2016; Mannan, Kamran, & Jeong, 2018; Jiang, Bian, & Tian, 2019). Until recently the successful removal of ocular artifact is mostly evaluated by visually inspecting the cleaned up EEG data and compare it with the non-cleaned data (Mannan, Kamran, & Jeong, 2018). Such an approach can easily lead to accidental removal of neural signals without being noticed because the amplitude of the artifacts can be as high as two orders of magnitude as large as the neural signals of interest. Recently (Sun, Chan, Hsiao, & Tang, 2020) provided a novel approach to quantitatively validate the extracted horizontal and vertical movement related ocular artifact and set an upper bound for any potential “contamination” of the ocular components by neural signals remained mixed with the ocular artifact. Specifically, the horizontal and vertical eye movement related components (H and V Comps) can be modeled as a pair of equivalent current dipoles (ECDs) with an ocular and non-neural origin accounting for over 95% variance in their scalp projections. Most interesting is the observation that the amplitudes of the recovered H and V Comps change systematically as a function of saccade directions and distances. Therefore, these components are more than mere artifacts but potentially containing signals indicating gaze positions. This result raises the possibility that these artifact components recovered from EEG data alone could be used as signals to predict gaze positions instead.

Here we explore this possibility by asking three questions: (1) how much variances in gaze positions can be explained by the H and V Comps; (2) how much data are needed to construct models to account for maximum amount of variance in gaze position; (3) how much data are needed to reach asymptotic performance in predicting gaze position using these modes. Specifically, we aim to demonstrate in a case study of a single participant that it is possible to construct models of gaze position using EEG alone and to use such models to predict gaze positions. To determine the optimal amount of data needed for producing asymptotic performance in model prediction, we studied one motivated participant performing the 8-direction and 2 distances directed saccadic eye movement task as same as the task in (Sun, Chan, Hsiao, & Tang, 2020) but with increased data length to 10 blocks. We established linear regression models with training dataset increased from 1 block to 9 blocks, and tested the performance (accuracy and reliability) on predicting gaze position on remaining dataset. The result showed that 4 bocks of data would also be sufficient to produce an asymptotic levels of prediction accuracy of 0.44 degrees of visual angles and reliability of 1.67 degrees of visual angles. Through this study, we introduce a novel concept of EEG based virtual eye tracking (EVET) and potentially enable the coregistration of eye movement and neuronal signal.

## 2. Materials and Methods

### 2.1 Participants

A single motivated normal adult served as the research participant to ensure the quality of behavioral data in this case study. All experimental procedures were conducted in accordance with the Human Research Ethics Committee of the University of Hong Kong and written informed consent was obtained from the participant prior to investigation.

### 2.2 Experimental Design

The present study has five components: (a) directed eye movement calibration task and simultaneous EEG and eye tracker data acquisition; (b) SOBI blind source separation of EEG data in increasing number of blocks (BKs) (BK_i_, i=1, 1-2, …1-10), resulting in 10 unmixing matrices _Wi_ (i=1, 2, …,10); (c) DANS automatic selection and validation of H and V ocular artifact components, performed on 10 sets of SOBI components generated by the 10 SOBI decompositions; (d) constructing 10 linear regression models (M_i_, i=1,2,…10) of gaze using the corresponding identified H and V Comps’ SRP amplitudes as predictor variables (derived from W_1_ through W_10_); (e) Using the model parameters of M_i_ to predict gaze position on data unused in model construction (BK_i+1 to 10_).

### 2.3 Behavioral task

The behavioral task is identical to that described in (Sun, Chan, Hsiao, & Tang, 2020) except that the length of the task was increased to five times of the original task (from 2 to 10 blocks of 16 trials). The participant was asked to sit in front of a computer screen in a quiet room and their head position was stabilized by a chinrest. The chinrest was adjusted such that participants’ eye level was aligned with the screen center with a viewing distance of 60 cm. The eye tracker was first calibrated using a 5-point eye-tracker calibration procedure repeatedly until the calibration error was below 0.5° of visual angle for both eyes. The directed eye movement task consisted of 160 directed eye movements from a central fixation point to a black dot presented on a white background, one at a time at 16 possible locations **(Fig. 1a)**, with 10 dots for each location. Among the 16 possible locations, 8 locations were 12.2° of visual angle away from the screen center and the other 8 locations were 6.1° of visual angle away from the screen center. These formed an inner (IN) and outer (OUT) ring of short versus long distance saccade target locations. In both visual angle conditions, the 8 locations were evenly distributed among all possible directions from the screen center (at 0°, 45°, 90°, 135°, 180°, 225°, 270°, and 315°, labeled as U, UR, R, DR, D, DL, L, UL respectively). Each dot was a circle subtending about 0.6° of visual angle (or a diameter of 16 pixels).

**Figure 1.**
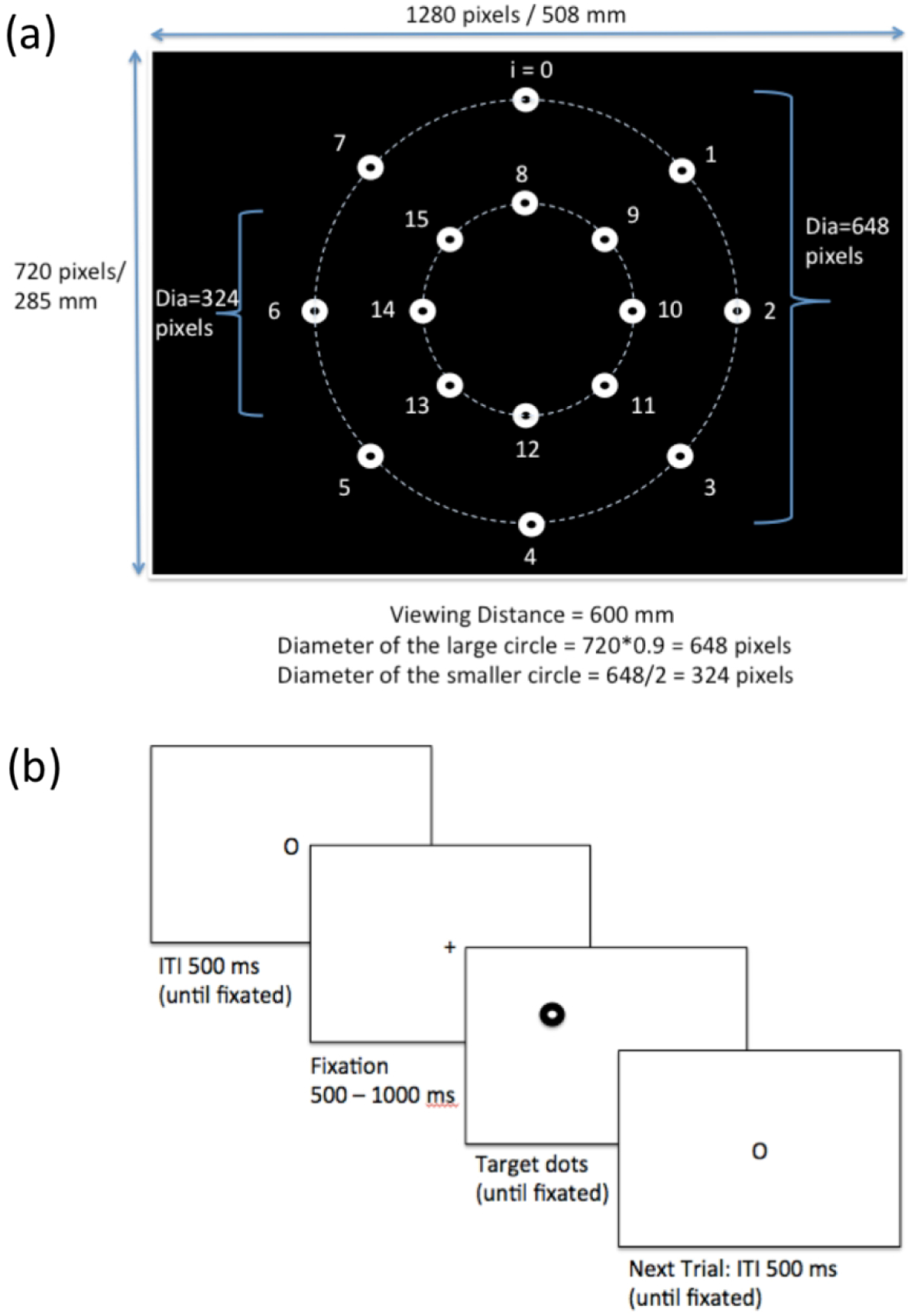
The directed saccadic eye movement task consisting of 10 blocks of 16 trials. **(a)**: arrangement of 16 target locations (8 directions and 2 distances). **(b)**: sequence of event within a trial. 149×204mm (221 × 221 DPI)

The participant was instructed to follow a sequence of dots presented at different locations in a random order one at a time **(Fig. 1b)**. Prior to each trial, the participant looked at the center of the screen where a fixation symbol “O” was displayed. A trial was initiated by the experimenter when the participant was fixated at the symbol “O”. The trial then began with a fixation cross “+” displayed at the center of the screen for a random duration between 500 ms and 1000 ms. Immediately after the offset of the fixation cross, a target dot appeared at one of the 16 possible locations randomly, to which the participant was instructed to move their eyes as fast and accurately as possible while avoiding blinks. Once a stable fixation on the target dot was detected by the eye tracker, then the dot disappeared and participants were allowed to blink until they saw the fixation symbol “O” again at the screen center for the next trial.

### 2.4 Eye Movement and EEG Data Acquisition and Processing

Eye movement data were continuously collected at a sampling rate of 60 Hz using the SMI REDn eye tracker, which was attached to a 17.5-inch monitor with a screen resolution of 1280×720 pixels. Only data from the dominant eye were used for eye-tracking data analysis. Trials in which the participant did not achieve valid fixations within 3 seconds were terminated by the experimenter and subsequently removed from data analysis. A valid fixation was defined by continuous eye gaze of at least 300 ms within a circular area around the target location with a radius of approximately 0.8° of visual angle. The number of trials excluded was 1.8 ± 0.56 (mean ± sem). The EEG data were recorded continuously along with the eye tracking data using the ANT EEGO ™Mylab system with 64 electrodes placed according to the standard system 10-20 at a sampling rate of 500Hz. E-prime 2.0 software communicated with the EEG system via a parallel port. Electrode impedance was kept below 20kΩ. The EEG data were notch-filtered (50hz) and high-pass filtered (0.1hz) before further processing.

### 2.5 SOBI Blind Source Separation

SOBI (Belouchrani, Abed-Meraim, Cardoso, & Moulines, 1997) is one of the BSS algorithms relatively less frequently used in analyzing EEG data. Detailed descriptions of SOBI’s usage (Tang, Pearlmutter, Malaszenko, & Phung, 2002; Tang, Pearlmutter, Malaszenko, Phung, & Reeb, 2002; Tang, Liu, & Sutherland, 2005; Sutherland & Tang, 2006; Tang, Sutherland, & Wang, 2006), validation (Tang, Sutherland, & McKinney, 2005; Lio & Boulinguez, 2013), and review of SOBI applications (Tang, 2010; Tang, Sutherland, & Yang, 2011; Urigüen & Garcia-Zapirain, 2015) can be found elsewhere. Here SOBI was applied to continuous EEG data collected during the entire 10 blocks of eye movement task to decompose the *n* - channel EEG data into *n* components (**Step 1, Fig. 2**), each of which corresponds to a recovered putative source that contributes to the scalp recorded EEG signals.

**Figure 2.**
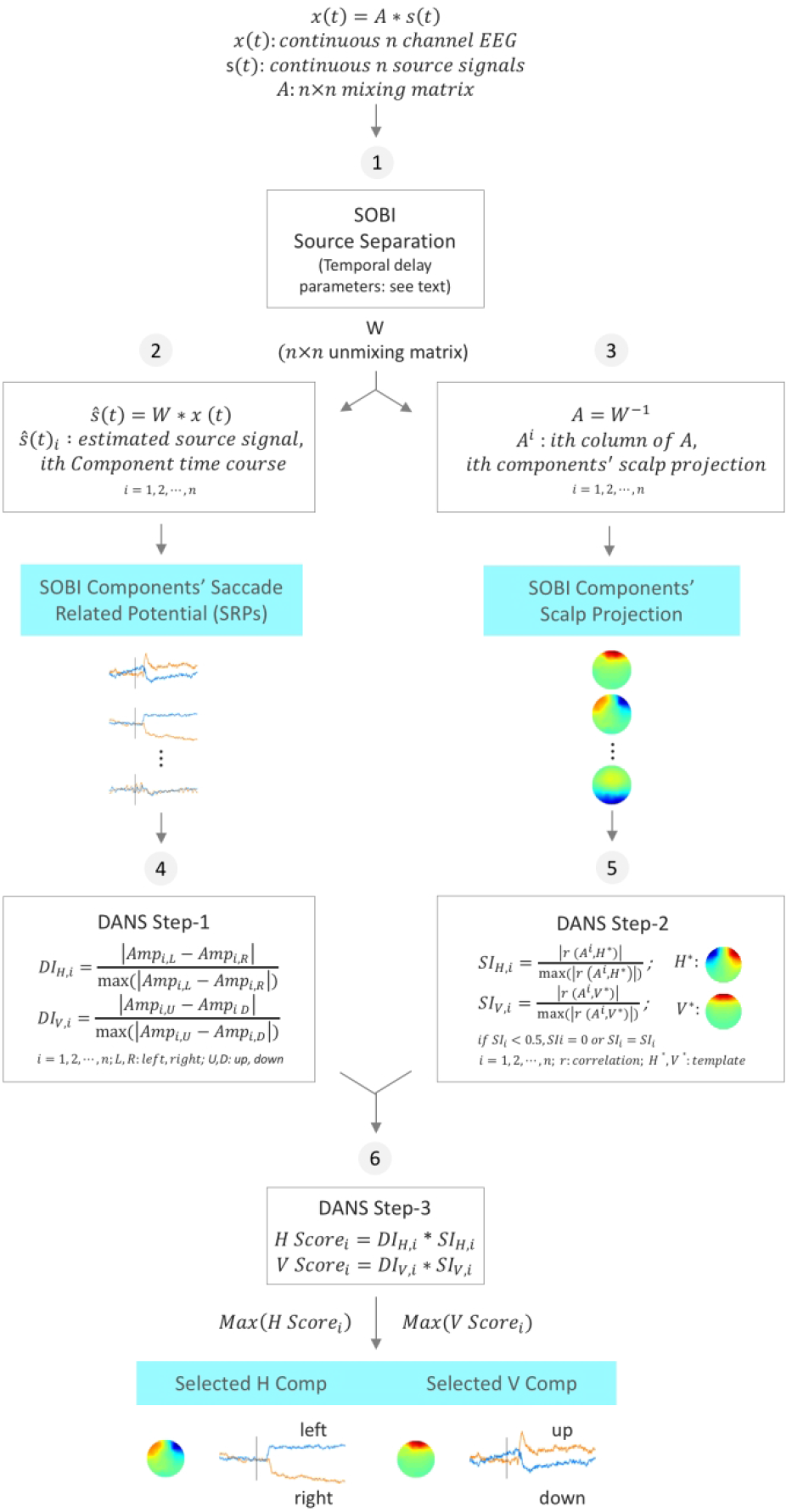
Flow chart of SOBI-DANS algorithm. (1) Generation of the unmixing matrix W; (2) temporal characterization of SOBI components via saccade-related potentials (SRPs); (3) spatial characterization of SOBI components via scalp projections; (4) computing the discriminant index (DI); (5) computing the similarity index (SI); (6) computing the H and V scores and selection of H and V Components. (Sun, Chan, Hsiao, & Tang, 2020) 280×510mm (112 × 112 DPI)

Let *x*(*t*) represent the *n* continuous time series from the *n* EEG channels, where *x*_*i*_(*t*) corresponds to the ith EEG channel. Because various underlying sources are summed via volume conduction to give rise to the scalp EEG, each of the *x*_*i*_(*t*) is assumed to be an instantaneous linear mixture of *n* unknown components or sources *s*_*i*_(*t*), via an unknown *n* × *n* mixing matrix *A*,

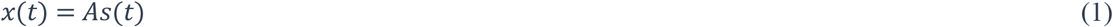

The putative sources, 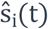 are given by

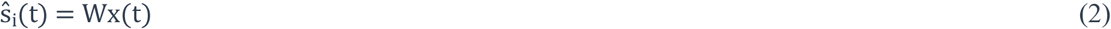

Where, *W* = *A*^−1^. SOBI finds the unmixing matrix *W* through an iterative process that minimizes the sum squared cross-correlations between one recovered component at time *t* and another at time *t* + *τ*, across a set of time delays. The following set of delays, *τs* (in ms), was chosen to cover a reasonably wide interval without extending beyond the support of the autocorrelation function:

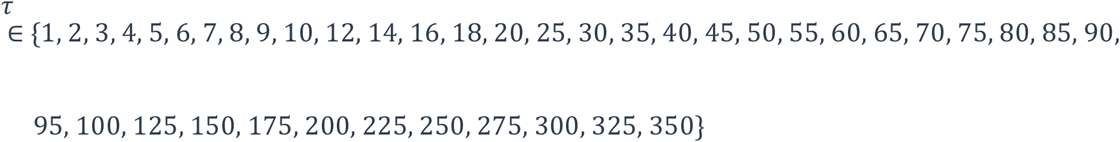

The *ith* component’s time course is given by 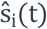, from which the SRP of each component can be generated (**Fig. 2 step 2**). The spatial location of the *i*th component is determined by the *i*th column of *A* (referred to as the component’s sensor weights or sensor space projection; **Fig 2, step 3**), where *A* = *W*^−1^.

### 2.6 DANS Identification of H and V Components

For each of the 10 SOBI decompositions using increasing blocks of data, the DANS algorithm was applied to all of SOBI components in order to select the best candidants for the ocular artifact components associated with horizontal and vertical eye movement. Therefore, a total of 10 pairs of H-V Comps were to be identified from the 10 SOBI decompositions (W1 through W10). The quality of the H-V Comps is expected to increase from with the increasing amount of EEG data used in deriving these ocular artifact components.

Each time, DANS generated two discriminant indices (DI_H_ and DI_V_) for every SOBI component to index temporal response selectivity (**Fig. 2 step 4**). DI_H_ and DI_V_ were defined as the normalized difference between two signed amplitudes of SRPs in response to the long saccades in opposite directions (DI_H_ : SRP_Left_-SRP_Right_; DI_V_ : SRP_Up_-SRP_Down_) in proportion to the maximum difference value (maximum DI=1.0). SRP amplitude was computed as the median amplitude within the 200-1200 ms after target onset and with a baseline correction window of 500 ms prior to target onset.

DANS also generated two similarity indices (SI_H_ and SI_V_) to index the resemblance of the component’s scalp projection with the known prototypical scalp projection maps of H and V ocular artifact components (**Fig. 2 Step 5**). SI_H_ and SI_V_ were defined as the normalized correlation between the scalp projection of a SOBI component and the projection of a prototypical H or V Comp, respectively (normalization by the maximum R, maximum = 1.0). A prototypical map can be found from any previously cited ICA and BSS studies of ocular artifact removal. If a SI is small than 0.5, then SI is set to zero, so that the corresponding component is effectively excluded from consideration since a component with SI < 50% of the SI_MAX_ is unlikely to be the single best candidate for being the component capturing the horizontal or vertical saccades related component. Finally, H and V scores, defined as the product of DI and SI, were computed for each component (Fig. 2 step 6). The components with the largest H or V scores were the final selected H or V Comps, respectively.

### 2.7 Systematic quantitative validation of H and V Comps

Each pair of the SOBI-DANS identified candidate H and V Comps were further validated via previously described validation procedure (Sun, Chan, Hsiao, & Tang, 2020) consists of (1) spatial validation via source localization using BESA (ECD modelling with one pair of symmetrically placed dipoles), (2) temporal validation via testing for a modulation of SRP amplitude as a function of saccade directions and distances.

#### 2.7.1 Spatial validation of H and V Components via equivalent current dipole (ECD) modeling

The purpose of spatial validation of the ocular artifact components is to ascertain to what extent these components capture and only capture signals of an ocular instead of a neural origin. The rationale is that if a component is truly an ocular component, then the estimated generators of the component’s scalp projection would not be located in neural tissues but near the physical apparatus that directly controls the eye balls and a pair of such dipoles should account for most of the variance in signal strength across the scalp projection map.

From Equation (1) and (2), the H and V Comps’ sensor space projection 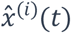 are given by

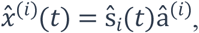

Where 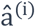 is a scalar given by the *i*th column of 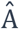(i.e., the component’s sensor weights) and 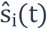 is in this case the average SRPs within a time window. A candidate’s average SRP following target onset can then be viewed as 3-D scalp voltage or current source density (CSD) maps. Using BESA (BESA 5.0, Brain Electrical Source Analysis; MEGIS Software, Munich, Germany), the scalp projection of the H and V Comps are modeled with a pair of symmetrically placed dipoles using a four-shell ellipsoidal model. The higher goodness of fit value an ECD model has for the H or V components, the greater confidence we have about the components’ ocular origin.

#### 2.7.2 Temporal validation via SRP amplitude tuning by saccade direction and distance

The rationale for temporal validation is that if the component’s time course reflects and only reflects signals associated with the movement of the eyes, then the SRP amplitude should vary systematically according to the direction and distance of eye movement. By testing for this prediction, we offer temporal validation of the ocular artifact components.

### 2.8 Modeling horizontal gaze position: model construction and prediction

To determine the amount of data needed for obtaining a model of gaze position with stable performance, we fitted 10 linear regressions with increasing block of data using the 10 pairs of H-V Comps extracted from 10 SOBI decompositions that generated 10 unmixing matrices. This process resulted in 10 sets of model parameters which were then used to make predictions of gaze positions from the SRP amplitude of H and V Comps for the remaining data that were unused for modeling construction (Fig. 3). Linear regressions were performed using General Linear Models function in SPSS (IBM SPSS 23, Armonk, NY: IBM Corp). The prediction error is computed as deviation of the predicted gaze position from the target position. The accuracy of model prediction is measured by the mean of errors and while the reliability of the model prediction is measured by the standard deviation of the errors. Three trials are missing due to missing eye-tracker confirmation of eye movement behavior (1.9%)

**Figure 3.**
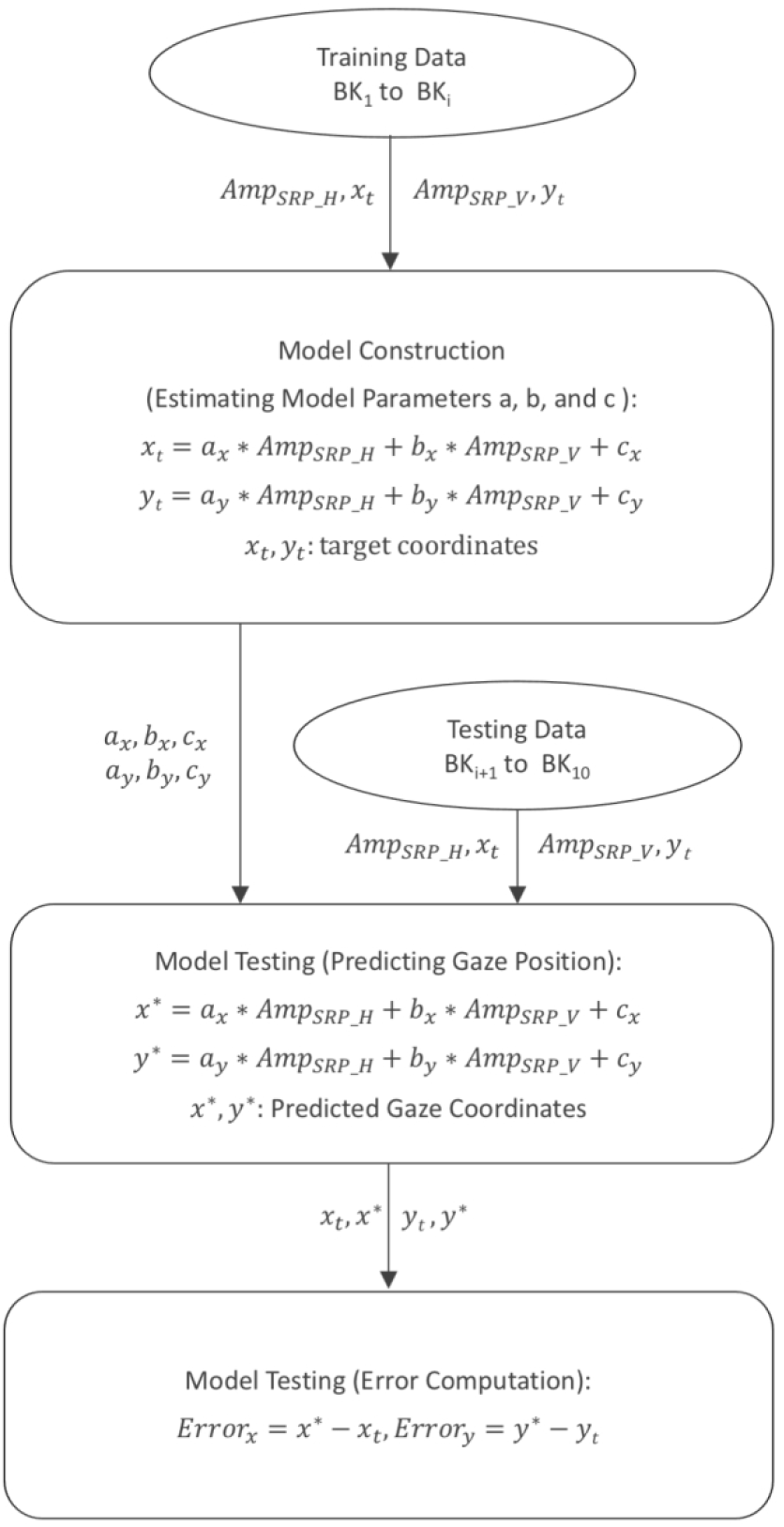
Flow chart of linear regression modeling of gaze position: (1) Model construction: training data from a subset of 10 blocks (BKs) of data used to estimate the 10 sets of model parameters a, b, and c; (2) Model testing: predicting gaze position using the estimated parameters and the SRP amplitude measures from the remaining data. 199×349mm (157 × 157 DPI)

## 3. Results

### 3.1 SOBI-DANS identification and validation of H and V comps

By applying SOBI-DANS method to EEG data of different length (1 to 10 blocks), 10 pairs of H and V Comps were extracted, identified, and validated temporally and spatially. **Fig. 4** offers an example (from M2, using two blocks of data) of such a pair, showing spatially the ECD models having a GoF of 97% and 98% respectively (**Fig. 4ab**) and temporally the SRP amplitudes being significantly modulated by saccade direction and amplitude (F(1, 24) = 52.231; p = 0.000; Partial η2 = 0.685) (**Fig. 4cd**). The significant correlations between response amplitudes and target gaze positions (**Fig. 4ef**, H Comp: R = 0.97; p = 0.000; N = 32 trials; V Comp: R = 0.79; p = 0.000; N = 32 trials), confirm that the two artifact components indeed contain great deal of information about gaze positions.

**Figure 4.**
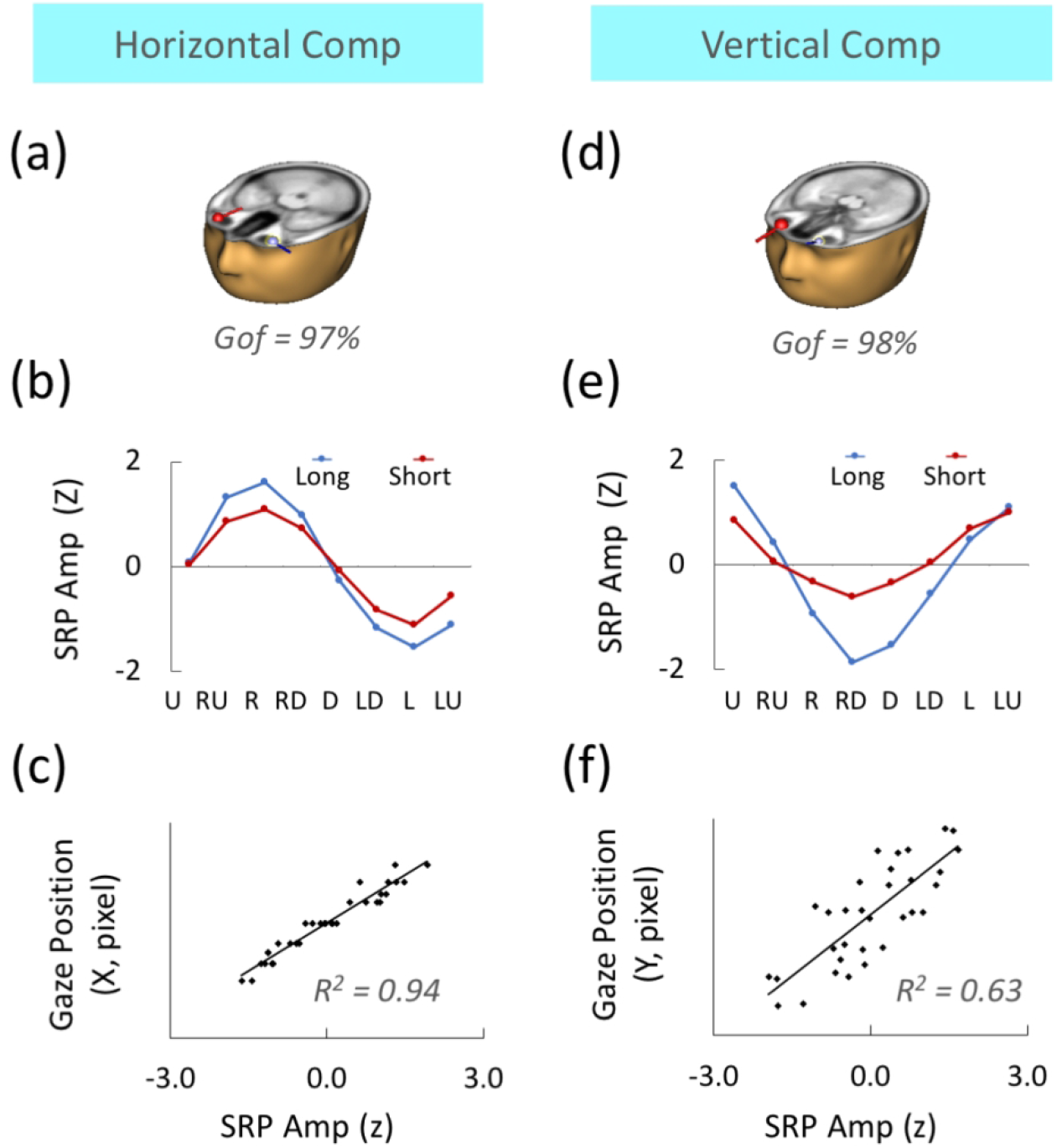
An example of SOBI-DANS extracted and identified horizontal and vertical eye movement related ocular artifact components (H and V Comps). Left: H Comp. Right: V Comp. **(ab)**: ECDs model goodness of fit and ECDs superimposed on sMRI. **(cd)** Tuning curves: amplitude of SRPs jointly modulated by saccadic eye movement direction and distance. **(ef)** Amplitude of SRPs are significantly correlated with gaze positions. H and V Comps extracted from the first two blocks of EEG data. 259×259mm (127 × 127 DPI)

The amount of variance in the *horizontal* dimension explained by the H and V Comps are particularly impressive reaching 94% (**Fig. 4e**). In contrast, the correlation for the V Comp and gaze position in the vertical dimension was 63%, only about 2/3 of that for H Comp (**Fig. 4f**). This relatively poor performance by SOBI on extracting component for vertical eye movement has been discussed in the past (Joyce, Gorodnitsky, & Kutas, 2004). Due to the scope of the present study, the remaining part of this paper will exclusively focus on the prediction of horizontal gaze position.

### 3.2 Modeling horizontal gaze position using SOBI-DANS recovered H and V Comps

We constructed 10 linear regression models of horizontal gaze position (Mi, i=1, 2, …10) with increasing amount of data (BKi, i= 1, 2, …, 10) and examined how the resulting model parameters vary as a function of the amount of data (BK) used by SOBI to generate the unmixing matrix Wi. **Fig. 5ab** shows the three parameters of the linear regression models reached a plateau with approximately 3 blocks of training data and more data do not make significant changes to the models. The goodness of the models as a function of the amount of training data can be indexed by the amount of variance in horizontal gaze position accounted for by the entire model **(Fig. 5c)** and by the model’s residual variance **(Fig. 5d)**. Fairly consistent with the trends in model parameters, model performance on the training data reached the plateau at three blocks of training data. This suggests that in order to obtain optimal model performance, at least four blocks data are needed. We also found that both the H and V Comps contribute significantly to the prediction of horizontal gaze position in the models with asymptotic performances (M1-M10). **Table 1** shows 2 estimated model parameters, their corresponding t statistics and p values. Note that while the parameters of H and V Comps (a_x_ and b_x_ respectively) are statistically significant in all models above M3, the contribution of H Comp is clearly bigger than V Comp, consistent with its larger t values in comparison to the V Comp’s t values. Also notice that the t values for H Comps are positive while the t values for V Comps are negative. This means the head of the participant was somewhat tilted to the right such that an upwards eye movement relative to the head produces an actual eye movement slightly to the right.

**Figure 5.**
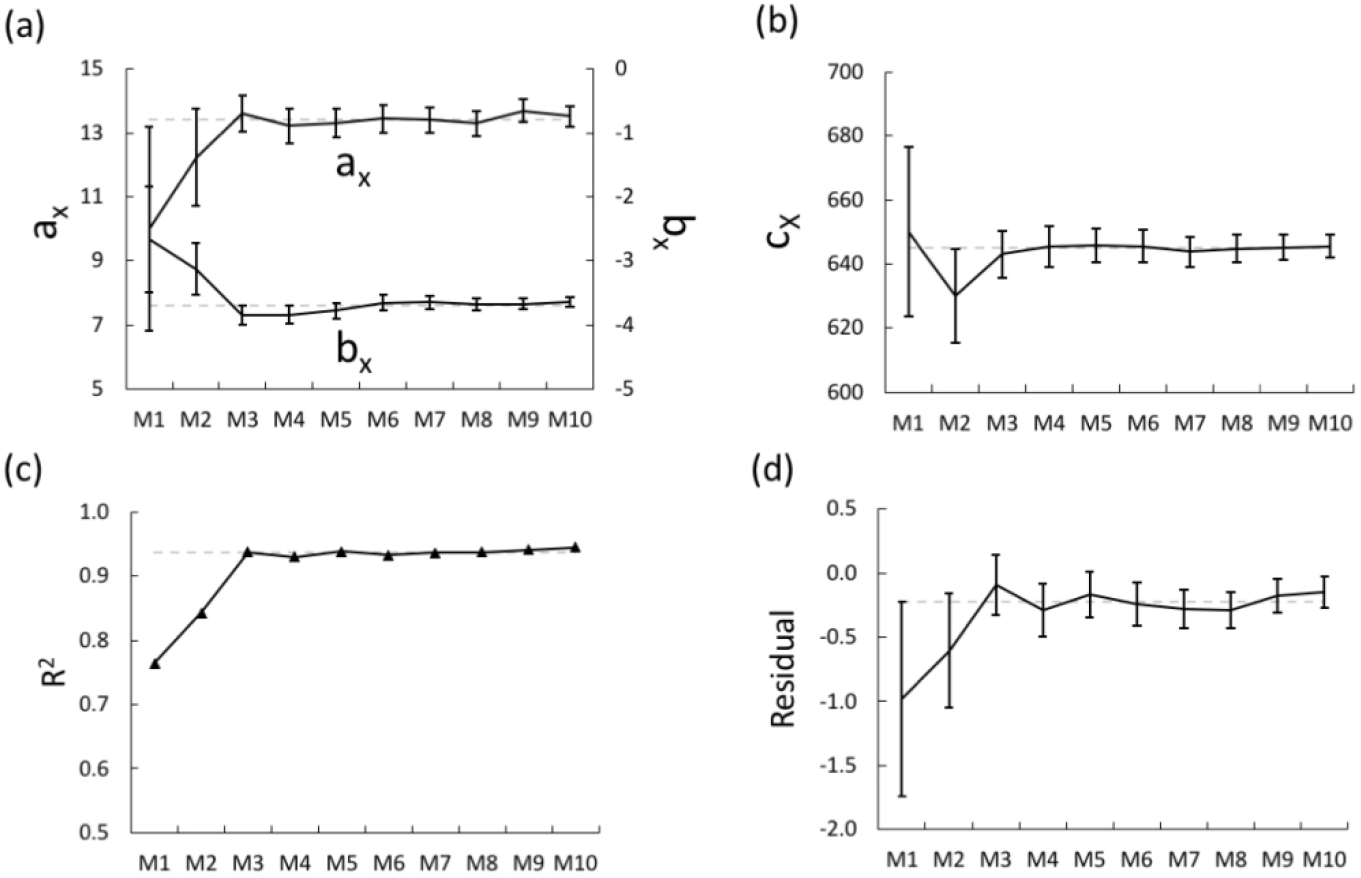
Parameters and performance of the linear regression models as a function of amount of data used (Mi: model constructed using data from block 1 to block i). **(a)** model coefficients for AmpSRP-H (ax) and AmpSRP-V (bx); **(b)** the constant cx; **(c)** R2: variance in gaze position (xt) accounted for by the model; **(d)** model residuals. 279×189mm (118 × 118 DPI)

**Table 1.** Contribution of H and V Comps for Predicting Horizontal Eye Movement across difference linear regression models (M_i_: the model constructed with the first i blocks of data)

| model training data<br>length (# of blocks) | $a_x$ | | $b_x$ | |
| --- | --- | --- | --- | --- |
|  | t | p | t | p |
| 1 | 5.720 | 0.000 | -1.432 | 0.176 |
| 2 | 10.443 | 0.000 | -1.890 | 0.069 |
| 3 | 22.389 | 0.000 | -2.520 | 0.015 |
| 4 | 24.625 | 0.000 | -3.550 | 0.001 |
| 5 | 30.404 | 0.000 | -4.029 | 0.000 |
| 6 | 32.670 | 0.000 | -3.863 | 0.000 |
| 7 | 37.082 | 0.000 | -4.349 | 0.000 |
| 8 | 39.427 | 0.000 | -4.704 | 0.000 |
| 9 | 45.441 | 0.000 | -4.428 | 0.000 |
| 10 | 48.790 | 0.000 | -5.196 | 0.000 |

### 3.3. Prediction of horizontal gaze position using SOBI-DANS recovered eye movement related components: model testing

We tested the first 9 models for prediction performance on the data not used in model construction (model i tested on block_i+1_ to block_10_). Predicted gaze positions were computed for M_1_to M_9_ by using the 9 sets of model parameters obtained in the previous section. Therefore, model i, Mi, generated from BK_1_ to BKi data, was tested on data from BK_i+1_ through BK_10_. Descriptive statistics are given in **Table 2.** Prediction errors as a function of the amount of data used in model construction are plotted both separately for each block (**Fig. 6ab**) and averaged across blocks (**Fig. 6cd**). Similar to the model construction curves shown in **Fig.** 5, the prediction errors change as a function of the length of data used in model construction with the accuracy (average) reached a stable level at M_4_ and the reliability (standard deviation) reached a stable level at M3. The final stable level of prediction performance was measured using prediction errors from M_4_ to M_9_ with the average model predictions of 0.44 ± 0.31 degrees of visual angle (**Fig. 6c**) and the standard deviation of model predictions of 1.67 ± 0.38 degrees of visual angle (**Fig. 6d**). Note that error bars on the curves (**Fig. 6cd)** indicating variations across different testing blocks. As an example, the 16 predicted gaze positions (from M5) are printed with the 16 target locations for one block data (BK9) (**Fig. 6e**, shaded line in **Table 2**).

**Figure 6.**
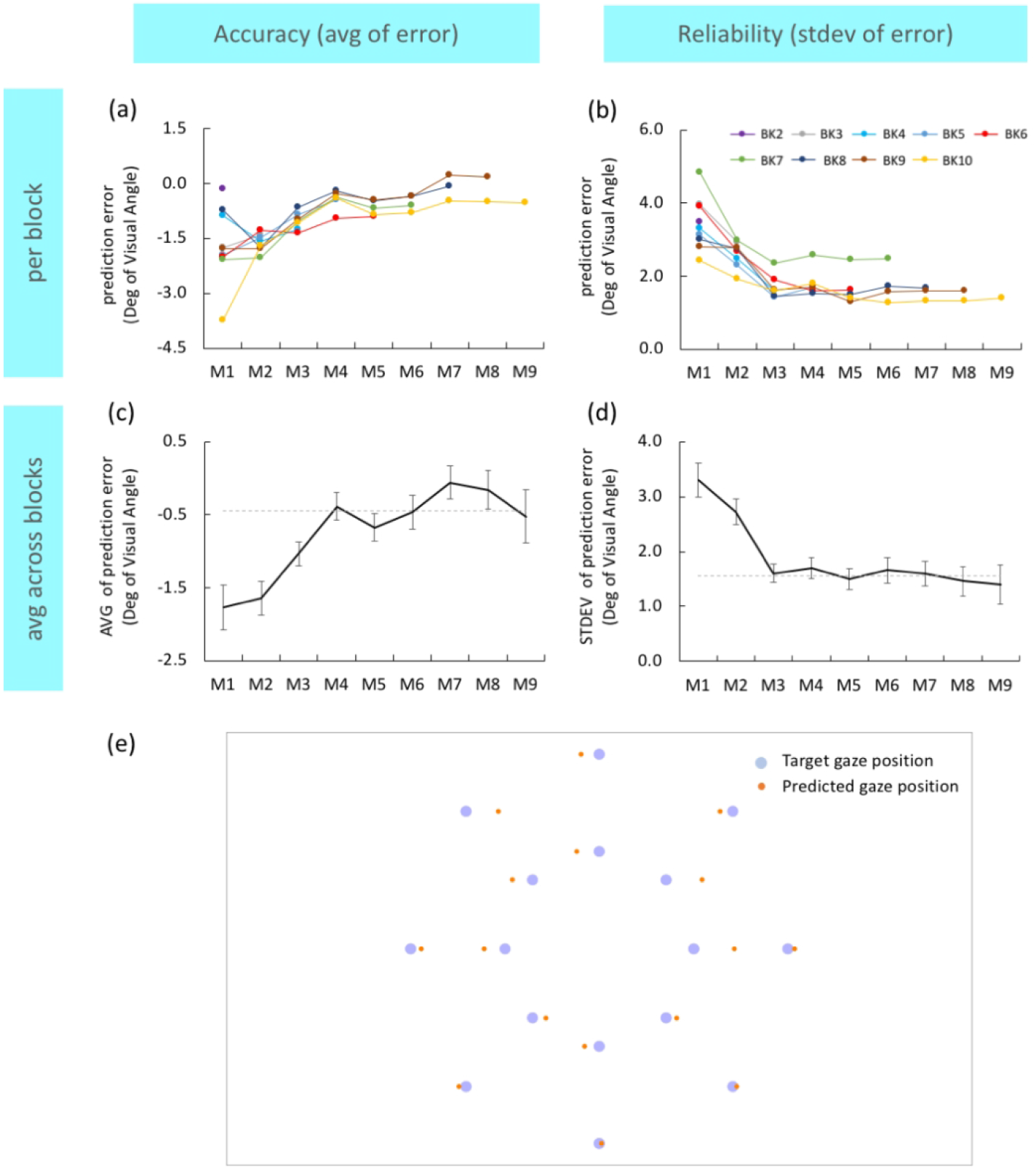
Prediction of horizontal gaze position x_p_ using remaining blocks of data. **(ac)** average prediction errors as a measure of model accuracy; **(bd)** standard deviation of prediction errors as a measure of model reliability. Prediction error as a function of amount of data used in model construction (M_i_, i=1, 2, …9), shown for each individual block of data **(ac)** and for all blocks of data pooled **(cd)**. **(e)** predicted gaze positions (from M5) and target locations in one block of data (BK9). 298×328mm (105 × 105 DPI)

**Table 2.**
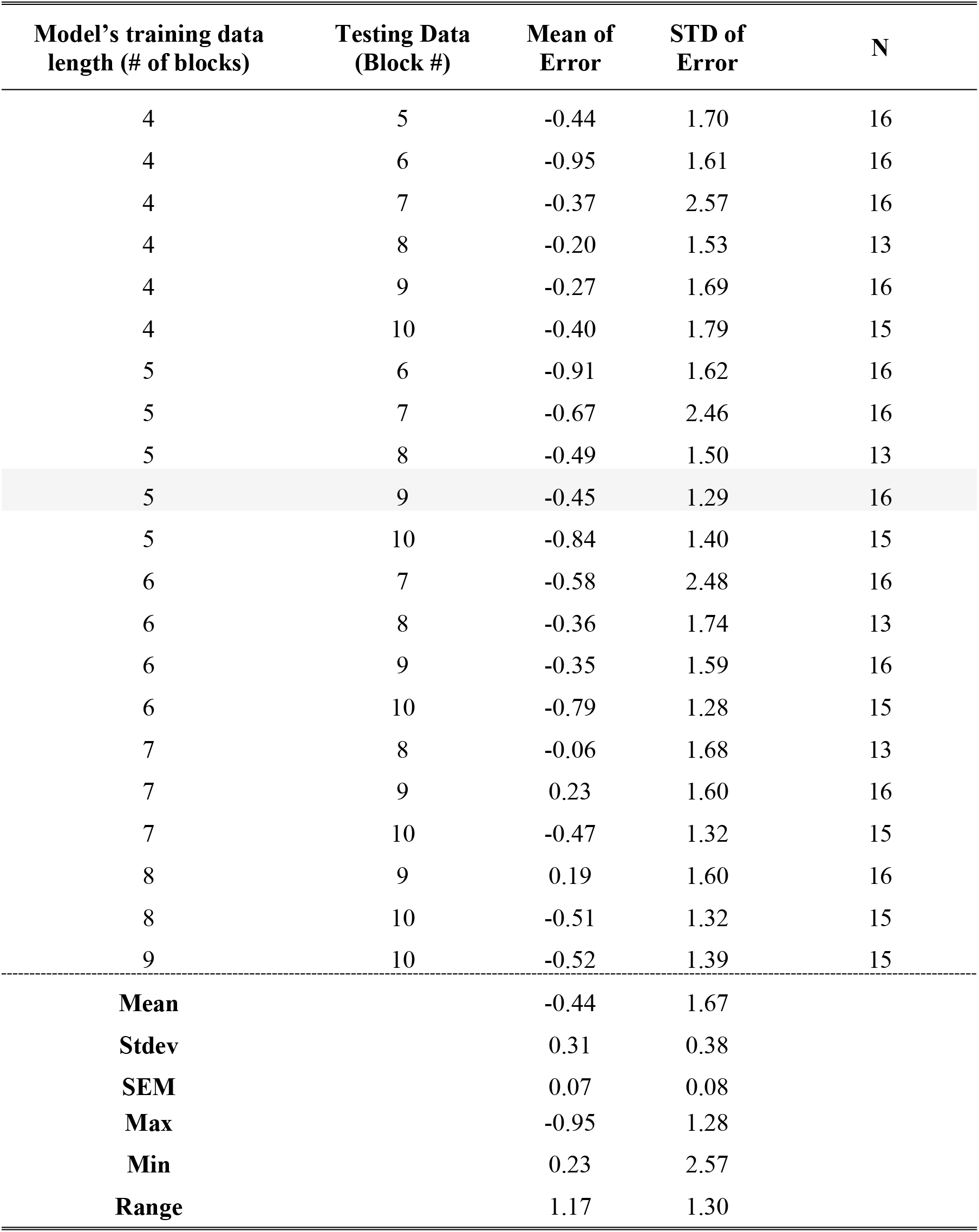
Mean and STD of prediction errors of the linear regression models (M_i_) for each combination of the model’s training data length and testing data block

| Model's training data length (# of blocks) | Testing Data (Block #) | Mean of Error | STD of Error | N |
| --- | --- | --- | --- | --- |
| 4 | 5 | -0.44 | 1.70 | 16 |
| 4 | 6 | -0.95 | 1.61 | 16 |
| 4 | 7 | -0.37 | 2.57 | 16 |
| 4 | 8 | -0.20 | 1.53 | 13 |
| 4 | 9 | -0.27 | 1.69 | 16 |
| 4 | 10 | -0.40 | 1.79 | 15 |
| 5 | 6 | -0.91 | 1.62 | 16 |
| 5 | 7 | -0.67 | 2.46 | 16 |
| 5 | 8 | -0.49 | 1.50 | 13 |
| 5 | 9 | -0.45 | 1.29 | 16 |
| 5 | 10 | -0.84 | 1.40 | 15 |
| 6 | 7 | -0.58 | 2.48 | 16 |
| 6 | 8 | -0.36 | 1.74 | 13 |
| 6 | 9 | -0.35 | 1.59 | 16 |
| 6 | 10 | -0.79 | 1.28 | 15 |
| 7 | 8 | -0.06 | 1.68 | 13 |
| 7 | 9 | 0.23 | 1.60 | 16 |
| 7 | 10 | -0.47 | 1.32 | 15 |
| 8 | 9 | 0.19 | 1.60 | 16 |
| 8 | 10 | -0.51 | 1.32 | 15 |
| 9 | 10 | -0.52 | 1.39 | 15 |
| **Mean** |  | -0.44 | 1.67 |  |
| **Stdev** |  | 0.31 | 0.38 |  |
| **SEM** |  | 0.07 | 0.08 |  |
| **Max** |  | -0.95 | 1.28 |  |
| **Min** |  | 0.23 | 2.57 |  |
| **Range** |  | 1.17 | 1.30 |  |

## 4. Discussion

We applied the previously described hybrid SOBI-DANS method to a single motivated individual performing 10 blocks of 16 directed eye movement task to demonstrate that it is possible to transform ocular artifacts into signals of gaze positions. We report an initial accuracy of ~ 0.44 degree of visual angle and reliability of ~ 1.67 degrees of visual angles, which are comparable to the precision of EOG-based eye tracking (Young & Sheena, 1975) and a combined EOG-EEG based eye tracking (Joyce, Gorodnitsky, King, & Kutas, 2002). This work offers the first prototype of a novel EEG-based virtual eye-tracker for computing horizontal gaze positions from EEG data alone, without having to place EOG electrodes on the face of the subject. New research directions for its potential applications and ways to further optimize the method are discussed.

### 4.1 Permitting free eye movement in EEG-based basic and clinical neuroscience research

Ocular artifacts associated with eye movement present a major problem in the cognitive neuroscience and clinical research. Partly as a solution to this problem, most visual paradigms used for experimental and diagnostic purposes require the eyes to be fixated at a particular location on a computer screen and then visual stimuli are presented to the fovea, which is assumed to be at the designated fixation point. A part of today’s understanding of neural mechanisms underlying visual cognition is based on this type of “passive” perception. Whether such an understanding can generalize from the context of passive to the more ecologically valid active perception remains a question until one can investigate neural mechanisms in the context of free eye movement.

The present finding points to a potential solution to this problem. Instead of minimizing the eye movement related artifacts by restricting eye movement and removing or throwing away these eye movement related signals as artifacts from the EEG data as documented in a number of review papers (Croft & Barry, 2000; Urigüen & Garcia-Zapirain, 2015; Islam, Rastegarnia, & Yang, 2016; Mannan, Kamran, & Jeong, 2018; Jiang, Bian, & Tian, 2019), we may now consider the option of making a directed saccadic eye movement task as an essential part of the experimental design and use the EEG data from this task to “calibrate” what may be viewed as a virtual EEG-based gaze tracker for predicting gaze positions in subsequent tasks that involve free eye movement. This option is feasible because we have previously shown that the eye movement related ocular artifact component can be decomposed from other neuronal components from EEG data collected during free viewing tasks (Tang, et al., 2006). Together these findings suggested that it is possible to concurrently collect the neural signals and EEG-derived gaze positions during cognitive tasks in the context of free eye movement.

### 4.2 Benefits and potential applications of an EEG based virtual eye tracker

If both neural signals and gaze positions can be measured concurrently all from EEG alone, then a separate device for eye tracking, the software and hardware for synchronizing the two devices, and the time delay problem in device synchronization could be by-passed. Therefore, the SOBI-DANS based EVET approach offers an inexpensive way to simultaneously track eye fixations and recording event-related brain potentials (ERPs) without the extra cost of an eye tracking hardware equipment. Furthermore, while near-infrared based eye trackers have the best resolution available, it requires the cameras to be able to see the eyes. This makes the study of eye movement behavior during sleep impossible. The SOBI-DANS based EVET approach would allow the by-passing of this problem.

These benefits bring out several potential applications of our new method and approach. The first is the tracking of the eyes during sleep to enable the investigation of patterns of eye movement during dreams and the relationship between eye movement patterns during the day and night (as was done in animal models by (Wilson & McNaughton, 1994) and to measure eye movement during memory reactivation, consolidation, and modification during sleep (Hu, et al., 2015). The second is tracking the gaze position during sentence reading, which was mostly done with the presentation of one word at a time while the subject maintains fixation (Kutas & Hillyard, 1980) with few exceptions (Dimigen, Sommer, Hohlfeld, Jacobs, & Kliegl, 2011). The SOBI-DANS based EVET approach would open up the possibility of a new protocol permitting natural reading of a horizontally printed sentences with free eye movement and potentially asking questions that could not be asked using the word-by-word presentation protocols, such as how words are represented across multiple brain regions during a saccade and between fixations. The third is tracking eye movement patterns and characteristics (including dwell times) in clinical neuroscience research, particularly when there is a clear need to measure both neural signals and gaze positions (Larrazabal, Garcia Cena, & Martinez, 2019).

### 4.3 Planning for EEG-based eye tracking

In general, application of the SOBI-DANS enabled EEG-based eye tracking consists of three steps-- first considering the spatial resolution of the eye tracking needed for solving your specific problems, second deciding on the calibration task that best covers the range of movement directions and distances, and third deciding on the amount of calibration data needed to build the optimal eye tracker for your specific application. If one is interested in identifying at a particular *moment* where the gaze position, the most relevant performance measure is the precision, i..e the standard deviation of the prediction error. Our current model has a precision equivalent to ~12 mm in the horizontal dimension. This translate into a potential error of about one word in English text printed at 12 point font at a reading distance of 40 cm. If one is interested in identifying over a *period of time* (as in dwell time), where in the visual scene or view the gaze has fallen, the more relevant performance is the accuracy measure, i.e. the average prediction error. The current model has an accuracy equivalent to 3 mm at the reading distance and 3 cm at 4 meters viewing distance.

### 4.4 Calibration of an EEG-based virtual eye-tracker

This paper presented a novel protocol for optimally constructing a model of gaze position from EEG data alone using SOBI-DANS. Although we used a particular calibration task consisting of 8 directions and 2 distances of saccadic eye movement, other calibration tasks may be used involving different spatial configurations or more/fewer target positions. Obviously, the choice should take into consideration of what the subsequent tasks of interest are and the spatial resolutions needed for measuring gaze position for the tasks of interest. If one only needs to judge which side of the left and right visual fields the subject is looking at, then you do not need to have a calibration task with densely distributed target locations. If one needs to determine which word in a sentence the subject has fixated on, then the distances between the target locations of your calibration task should be as small as the shortest saccade length in reading the sentence.

Without prior knowledge, as a conservative decision, the calibration task used in this present study consisted of 10 repetitions of each target location based on the assumption that 10 repetitions of 16 target locations (~15 min of EEG data) should be sufficient for reaching the optimal extraction of the movement related ocular artifacts by SOBI-DANS. The model construction and prediction curves have three regions: the rapid change region in which the model parameters and its prediction errors continue to change with increasing amount of training data (here BK1-3); the final stable region in which an increase in the amount of data does not produce further changes in model parameters or predictions errors (here BK6-10); the region in-between in which model parameters and predictions still show small fluctuations from one block to another (here BK3-6) (**Fig. 5 and Fig. 6)**. If one needs the highest precision and if it is practical to run a longer calibration task, one could set the calibration data length to be the smallest number of blocks in the final stable region (6 blocks). If the tasks of interest do not require the highest precision or if a longer calibration task is not practical, then the smallest number of blocks in the middle region could be chosen (4 blocks). One should avoid the rapid change region (1-2 blocks) in setting the calibration data length.

### 4.5 Future work

One of the promising directions for performance improvement is the consideration of other types of models, such as neural networks and support vector machines, which are known to have generally superior performance than linear regression models for non-linear functions. Additionally, current work has focused on solving the easier problem of predicting horizontal gaze position and the harder problem of predicting vertical gaze positions (Joyce et al 2002) remains to be explored. In summary, we hope that the technical capability afforded by this new concept of virtual eye tracker will enable new areas of investigation, such as neurophysiological investigation of natural reading, learning and memory during sleep, better devices for neurofeedback control, and clinical neuroscience problems. Perhaps, in the future, installing a software would be sufficient to simultaneously collect neuronal signals and track eye movements.

## Acknowledgement

This work is supported by the University of Hong Kong grant [grant number 104004683] and the Sweeting Memorial Fund to AC Tang; the Research Grant Council [grant number 17609117] to Hsiao.

## Notes

### Competing Interest Statement

The authors have declared no competing interest.

